# Metformin improves glycemia independently of skeletal muscle AMPK via enhanced intestinal glucose clearance

**DOI:** 10.1101/2022.05.22.492936

**Authors:** Rasmus Kjøbsted, Jonas M. Kristensen, Jesper B. Birk, Nicolas O. Eskesen, Kohei Kido, Nicoline R. Andersen, Jeppe K. Larsen, Marc Foretz, Benoit Viollet, Flemming Nielsen, Kim Brøsen, Niels Jessen, Ylva Hellsten, Kurt Højlund, Jørgen F.P. Wojtaszewski

**Affiliations:** August Krogh Section for Molecular Physiology, Department of Nutrition, Exercise and Sports, Faculty of Science, University of Copenhagen, Copenhagen, DK-2100, Denmark; Université Paris Cité, CNRS, INSERM, Institut Cochin, F-75014 Paris, France; Clinical Pharmacology and Pharmacy, Department of Public Health, University of Southern Denmark, Odense, DK-5000, Denmark; Steno Diabetes Center Aarhus, Aarhus University Hospital, Aarhus, DK-8200, Denmark; August Krogh Section for Human Physiology, Department of Nutrition, Exercise and Sports, Faculty of Science, University of Copenhagen, Copenhagen, DK-2100, Denmark; Steno Diabetes Center Odense, Odense University Hospital, Odense, DK-5000, Denmark; Department of Clinical Research & Development of Molecular Medicine, University of Southern Denmark, Odense, DK-5000, Denmark

**Author notes:** **Correspondence:** (R.K.), (J.F.P.W.).

## Abstract

Metformin is an inexpensive oral anti-hyperglycemic agent used worldwide as a first-choice drug for the prevention of type 2 diabetes mellitus (T2DM). Although current view suggests that metformin exerts its anti-hyperglycemic effect by lowering hepatic glucose production, it has been proposed that metformin also reduce hyperglycemia by increasing glucose uptake in skeletal muscle via activation of AMP-activated protein kinase (AMPK). Herein, we demonstrate in lean and diet-induced obese (DIO) male and female mouse models that the anti-hyperglycemic effect of metformin occurs independently of muscle AMPK, and instead relies on elevated intestinal glucose clearance. Furthermore, we report that the AMPK activity is elevated in skeletal muscle from patients with T2DM following chronic metformin treatment, but this is not associated with enhanced peripheral insulin sensitivity. These results argue against existing paradigms and emphasize the non-essential role of muscle AMPK but important role of the intestine for the anti-hyperglycemic effect of metformin.

## Introduction

Metformin has been used worldwide as a first-choice drug to treat type 2 diabetes mellitus (T2DM) due to its blood glucose-lowering properties and high safety profile (Bailey and Turner, 1996). Several attempts have been made to elucidate the exact mechanism(s) by which metformin exerts its effect on blood glucose levels (Foretz et al., 2019; Rena et al., 2017). Some evidence points towards a role of metformin action to suppress hepatic glucose production (Foretz et al., 2010; Fullerton et al., 2013; Hunter et al., 2018; Madiraju et al., 2014, 2018; Miller et al., 2013; Shaw et al., 2005) but other sites of metformin action have also been proposed including the intestine (Bailey et al., 2008; Koffert et al., 2017; Ma et al., 2022; Rittig et al., 2021), gut microbiota (Wu et al., 2017), gut-brain-liver neuronal axis (Duca et al., 2015), and skeletal muscle (Musi et al., 2002; Zhou et al., 2001). This highlights that metformin likely exploits various tissue-specific mechanisms to regulate blood glucose levels.

Results from *in vitro* studies using supra clinical metformin concentrations suggest that metformin exerts some of its glucose-lowering properties by selective inhibition of mitochondrial complex I activity (Brunmair et al., 2004; El-Mir et al., 2000; Owen et al., 2000), which leads to a decrease in ATP production and a subsequent increase in the AMP/ATP ratio (Foretz et al., 2010; Stephenne et al., 2011). The crucial cellular energy sensor AMP-activated protein kinase (AMPK) functions to maintain energy homeostasis and is activated by energy stress that perturbs the intracellular AMP/ATP and/or ADP/ATP ratio (Lin and Hardie, 2018). When activated in skeletal muscle, AMPK promotes glucose uptake and fatty acid oxidation while suppressing protein synthesis (Garcia and Shaw, 2017; Kjøbsted et al., 2018). In a seminal study from 2001, metformin was reported to activate AMPK in isolated rat skeletal muscle. This occurred concomitant with an increase in glucose uptake that was additive to the effect of insulin (Zhou et al., 2001). Based on these observations, it has been hypothesized that metformin may function to lower blood glucose levels *in vivo* via inhibition of mitochondrial complex I activity in skeletal muscle that activates AMPK and thus promotes glucose uptake (Cokorinos et al., 2017) and enhance insulin sensitivity (Kjøbsted et al., 2015).

Here, we sought to test the hypothesis that AMPK mediates metformin action by augmenting glucose uptake and insulin sensitivity in skeletal muscle. By examining the consequences of conventional and conditional muscle-specific knockout of AMPKα1/α2 in metformin-treated lean and diet-induced obese (DIO) male and female mouse models, we report that lack of AMPK activity in skeletal muscle does not compromise the ability of metformin to lower blood glucose levels or improve whole-body glucose tolerance. Furthermore, we find elevated AMPK activity in skeletal muscle from metformin-treated patients with T2DM, which is not associated with enhanced peripheral insulin sensitivity. In contrast, our study provides evidence to support that the anti-hyperglycemic effect of metformin involves enhanced intestinal glucose clearance.

## Results

### Muscle AMPK is dispensable for the acute blood glucose lowering effect of metformin

Prior to testing the effect of metformin in AMPK-deficient mice, we sought to confirm the effect of a single dose of metformin (250 mg kg^-1^) commonly used to elicit a blood glucose-lowering effect in mice (Foretz et al., 2010; Fullerton et al., 2013; Hunter et al., 2018; Miller et al., 2013). Compared to the vehicle-treated control groups, acute intraperitoneally (i.p.) administered metformin decreased blood glucose levels by ~20-30% after 60 min to a similar extent in lean and diet-induced obese (DIO) wildtype (WT) mice (Figures 1A-1C, S1A and S1B). This occurred despite that plasma, liver, and muscle metformin concentrations were several fold higher in DIO vs. lean mice (Figure 1D). Metformin increased the AMP/ATP ratio in the liver but this did not translate into elevated liver AMPK activity/signaling (Figures S1C-1F). Metformin did not increase either the AMP/ATP ratio or AMPK activity/signaling in skeletal muscle (Figures 1E-G and S1G). Nor did metformin significantly increase plasma insulin levels or muscle and liver insulin signaling (Figures S1H and S1I). This suggests that lean and DIO mice are equally susceptible to the acute anti-hyperglycemic effect of metformin that occurs in the absence of elevated AMPK activity in muscle and liver.

**Figure 1.**
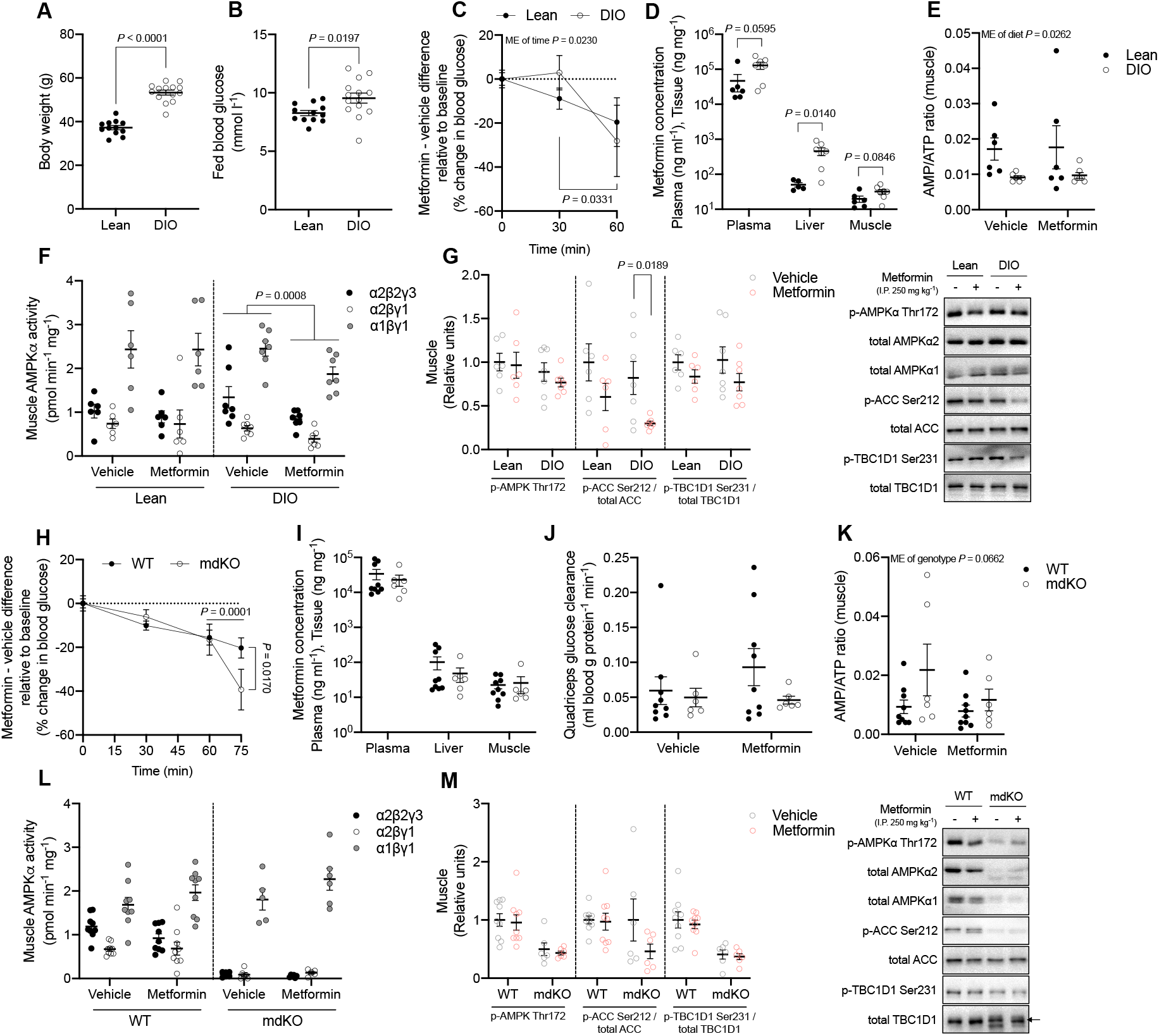
Acute metformin treatment lowers blood glucose levels in lean and DIO mice independently of enhanced AMPK activity in skeletal muscle. (A and B) Body weight (A) and fed blood glucose (B) in lean and DIO mice measured the morning after a night of free access to respective diets (n = 12/14, Lean/DIO). (C) Effect of i.p. administered metformin (250 mg kg^-1^) vs. vehicle on blood glucose levels (%) in lean and DIO mice (n = 6/7, Lean/DIO). (D) Metformin concentrations in plasma (ng ml^-1^) as well as liver and quadriceps muscle (ng mg^-1^) 60 min after dosing (n = 5-7 per group). (E-G) AMP/ATP ratio (E), AMPK activity of the α2βγ1, α1βγ1 and α2β2γ3 complexes (F) and phosphorylation of AMPKα Thr172, ACC Ser212, and TBC1D1 Ser231 (G) in quadriceps muscle of vehicle- and metformin-treated mice (n = 6-7 per group). Right: representative immunoblots. (H) Effect of i.p. administered metformin (250 mg kg^-1^) vs. vehicle on blood glucose levels (%) in fed and lean conventional muscle-specific AMPKα1α2 double KO female mice (mdKO) and wildtype (WT) littermates (n = 6-11 per group) (I) Metformin concentrations in plasma (ng ml^-1^) as well as liver and quadriceps muscle (ng mg^-1^) from AMPK mdKO mice and WT littermates 75 min after dosing (n = 6/9, mdKO/WT). (J) Glucose clearance in quadriceps muscle from AMPK mdKO mice and WT littermates determined by the use of ^3^H-2-deoxy-D-glucose tracing (n = 6/9, mdKO/WT). (K) AMP/ATP ratio in quadriceps muscle. (L and M) AMPK activity of the α2βγ1, α1βγ1 and α2β2γ3 complexes (L) and phosphorylation of AMPKα Thr172, ACC Ser212, and TBC1D1 Ser231 (M) in quadriceps muscle of vehicle- and metformin-treated AMPK mdKO mice and WT littermates (n = 6/9, mdKO/WT). Right: representative immunoblots. Data represent mean ± SEM. Data in (A, B, D, G, I, and M) were analyzed using a homoscedastic two-tailed student’s t-test. Data in (C, E, F, H, J, K and L) were analyzed using a two-way ANOVA with (C and H) or without (E, F, J, K and L) repeated measures and Sídák post hoc analyses for multiple comparisons. HFD, high-fat diet. ME, main effect.

To further investigate the role of muscle AMPK for the anti-hyperglycemic effect of metformin, lean muscle-specific AMPKα1α2 double knockout (mdKO) mice and WT littermates matched on weight, body composition and blood glucose levels (Figures S1J-L) were administered a single dose of metformin (i.p. 250 mg kg^-1^). Compared to the vehicle-treated control mice, acute metformin treatment lowered blood glucose levels by ~20-40% after 75 min regardless of muscle AMPK deficiency (Figures 1H, S1M and S1N). Importantly, metformin levels in plasma, liver, and muscle were comparable between mdKO and WT mice (Figure 1I). Employing a ^3^H-2-deoxy-D-glucose tracer in combination with the acute metformin treatment permitted the assessment of tissue-specific glucose uptake. We observed that the glucose clearance, AMP/ATP ratio as well as AMPK activity/signaling were not increased in muscle from mdKO or WT mice in response to metformin (Figures 1J-M, S1O and S1P). Notably, liver, but not muscle, glycogen levels decreased by ~50% in response to metformin (Figures S1Q and S1R). Plasma insulin levels as well as muscle insulin signaling were unaffected by metformin (Figures S1S and S1T) and therefore cannot explain the observed decrease in blood glucose levels. These findings demonstrate that the acute anti-hyperglycemic effect of i.p. administered metformin occurs independently of enhanced AMPK activity and glucose uptake in skeletal muscle.

### Acute metformin treatment lowers blood glucose levels by enhancing intestinal glucose clearance

Since metformin is commonly administered orally, we sought to rule out potential bias of i.p. administered metformin that may conceal a relevant role of muscle AMPK for the anti-hyperglycemic effect of metformin. Compared to the vehicle-treated control mice, a single dose of metformin administered by oral gavage (p.o. 250 mg kg^-1^) lowered blood glucose levels by ~40-50% in mdKO and WT mice (Figures 2A, S2A and S2B). Metformin levels in plasma, liver, and muscle were similar between mdKO and WT mice following oral dosing (Figure 2B) and comparable to findings after i.p. administration (Figure 1D and I). Glucose clearance increased slightly in brain tissue but was otherwise unaffected in muscle, adipose, and heart tissue following oral metformin dosing (Figures 2C and S2C). The AMP/ATP ratio and AMPK activity/signaling in muscle were also unaffected by orally administered metformin (Figures 2D-F and S2D). Once again, we observed that metformin decreased liver, but not muscle, glycogen levels (Figures S2E and S2F). Plasma glucagon levels were increased following oral and intraperitoneal administration of metformin (Figure S2G) that may explain the metformin-induced decrease in liver glycogen. Neither plasma insulin levels nor muscle insulin signaling was affected by orally administered metformin (Figures S2H and S2I).

**Figure 2.**
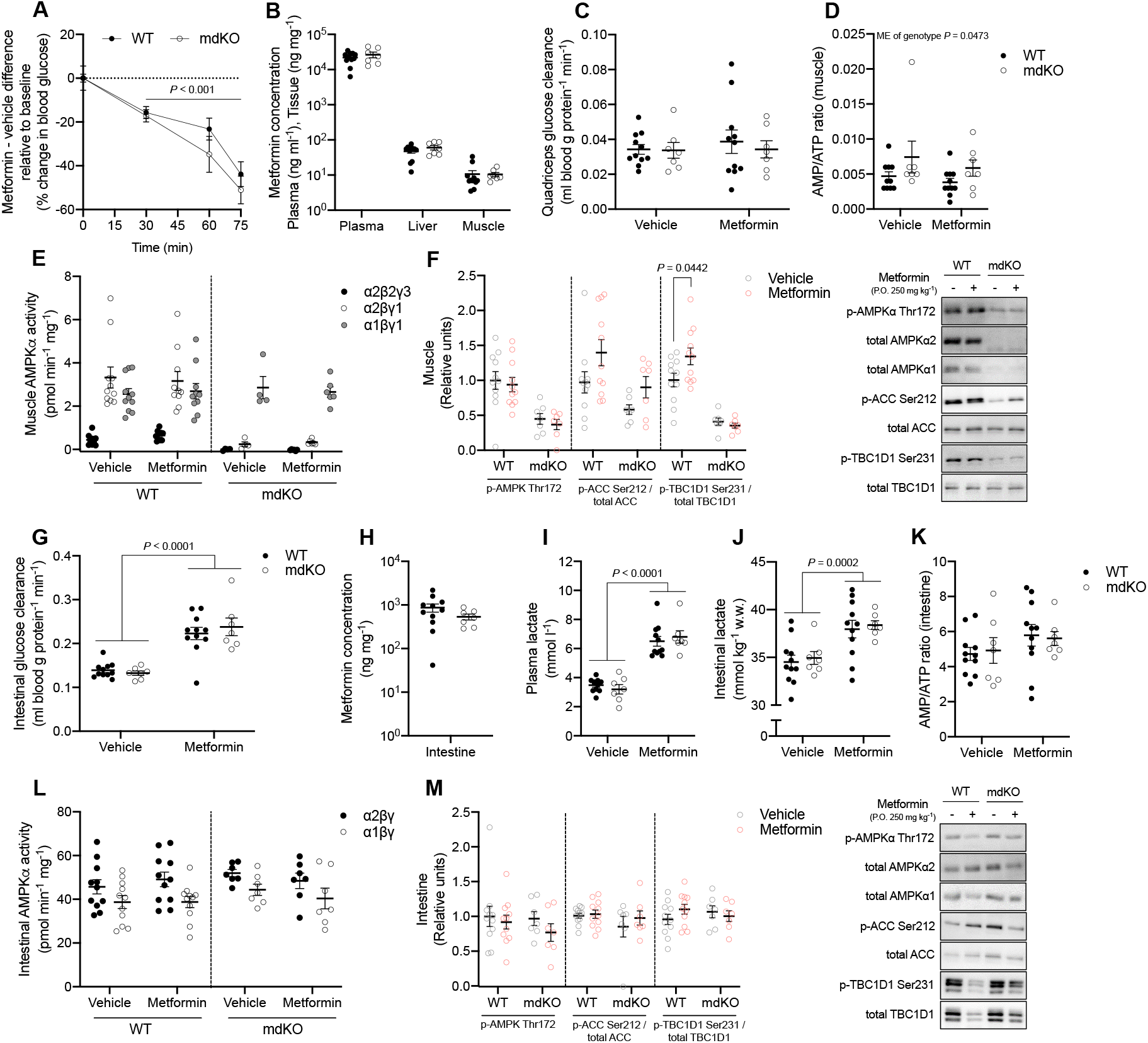
Acute metformin treatment lowers blood glucose levels by enhancing intestinal glucose clearance. (A) Effect of orally dosed metformin (250 mg kg^-1^) vs. vehicle on blood glucose levels (%) in lean AMPK mdKO mice and WT littermates. (B) Metformin concentrations in plasma (ng ml^-1^) as well as liver and quadriceps muscle (ng mg^-1^) 75 min after dosing. (C) Glucose clearance in quadriceps muscle following vehicle/metformin treatment determined by the use of ^3^H-2-deoxy-D-glucose tracing (n = 7/11, mdKO/WT). (D) AMP/ATP ratio in quadriceps muscle (E and F) AMPK activity of the α2βγ1, α1βγ1 and α2β2γ3 complexes (E) and phosphorylation of AMPKα Thr172, ACC Ser212, and TBC1D1 Ser231 (F) in quadriceps muscle of vehicle- and metformin-treated lean muscle-specific AMPKa1a2 KO mice and WT littermates. Right: representative immunoblots (G) Intestinal glucose clearance of orally metformin-dosed lean AMPK mdKO mice and WT littermates determined by the use of ^3^H-2-deoxy-D-glucose tracing (n = 7/11, mdKO/WT). (H) Metformin concentration in intestinal tissue (ng mg^-1^). (I) Lactate levels in intestinal tissue of orally metformin-dosed lean AMPK mdKO mice and WT littermates. (J) Plasma lactate concentration 75 min after dosing. (K-M) AMP/ATP ratio (K), AMPKα2 and AMPKα1 complex-specific activity (L) and phosphorylation of AMPKα Thr172, ACC Ser212, and TBC1D1 Ser231 (M) in the intestine after orally dosed metformin. Right: representative immunoblots Data represent mean ± SEM. Data in (B, F, H, and M) were analyzed using a homoscedastic two-tailed student’s t-test. Data in (A, C, D, E, G, and I-L) were analyzed using a two-way ANOVA with (A) or without (C, D, E, G, and I-L) repeated measures and Sídák post hoc analyses for multiple comparisons. w.w, wet weight. ME, main effect.

Several lines of evidence indicate that the gastrointestinal tract is involved in the acute anti-hyperglycemic effect of metformin (Bailey et al., 2008; Koffert et al., 2017; Ma et al., 2022; Rittig et al., 2021). In accordance, we observed that metformin increased glucose clearance by ~75% in the proximal part (duodenum and jejunum) of the intestine (Figure 2G) that displayed ~10- and 70-fold higher metformin concentrations compared to the liver and muscle, respectively (Figure 2H). Following uptake into the intestine, glucose is directed through glycolysis to form lactate since metformin increases anaerobic glucose metabolism in the intestine (Bailey et al., 2008; Rittig et al., 2021). This likely contributes to increased blood lactate levels that are associated with regular metformin use (Bailey and Turner, 1996). In line, we observed that metformin increased plasma lactate levels (Figure 2I) as well as lactate levels in the intestine, but not in liver and muscle tissue (Figures 2J and S2J). We did not find an elevated AMP/ATP ratio or increased AMPK activity/signaling in the intestine following metformin treatment (Figures 2K-M and S2K). This supports our recent findings showing an intact glucose-lowering effect of acute metformin treatment in intestinal-specific AMPKα1α2 knockout mice (Olivier et al., 2021).

### Acute metformin treatment enhances whole body glucose tolerance independently of muscle AMPK

We have previously reported that prior exercise-, contraction- and AICAR-induced activation of AMPK potentiates insulin action in skeletal muscle (Kjøbsted et al., 2015, 2017). Therefore, we sought to investigate whether muscle AMPK deficiency would diminish the effect of prior acute metformin treatment to promote a faster disappearance of blood glucose during a glucose tolerance test. We found that the glucose-lowering effect of prior metformin administration (i.p. and p.o. 250 mg kg^-1^) was preserved during a glucose tolerance test in conventional and conditional muscle-specific lean and DIO AMPKα1α2 KO mice (Figures 3A-C and S3A-S). Plasma insulin levels were similar between WT and KO mice but slightly higher in metformin-vs. vehicle-treated mice (Figures 3D and S3T-V). This demonstrates that the enhancing effect of metformin on whole-body glucose tolerance occurs independently of muscle AMPK but may in part involve enhanced secretion of insulin. The latter is supported by previous studies in human and animal models and likely triggered by an augmented secretion of glucagon-like peptide-1 (GLP-1) from the enteroendocrine L cells (Maida et al., 2011; Mulherin et al., 2011; Vardarli et al., 2014).

**Figure 3.**
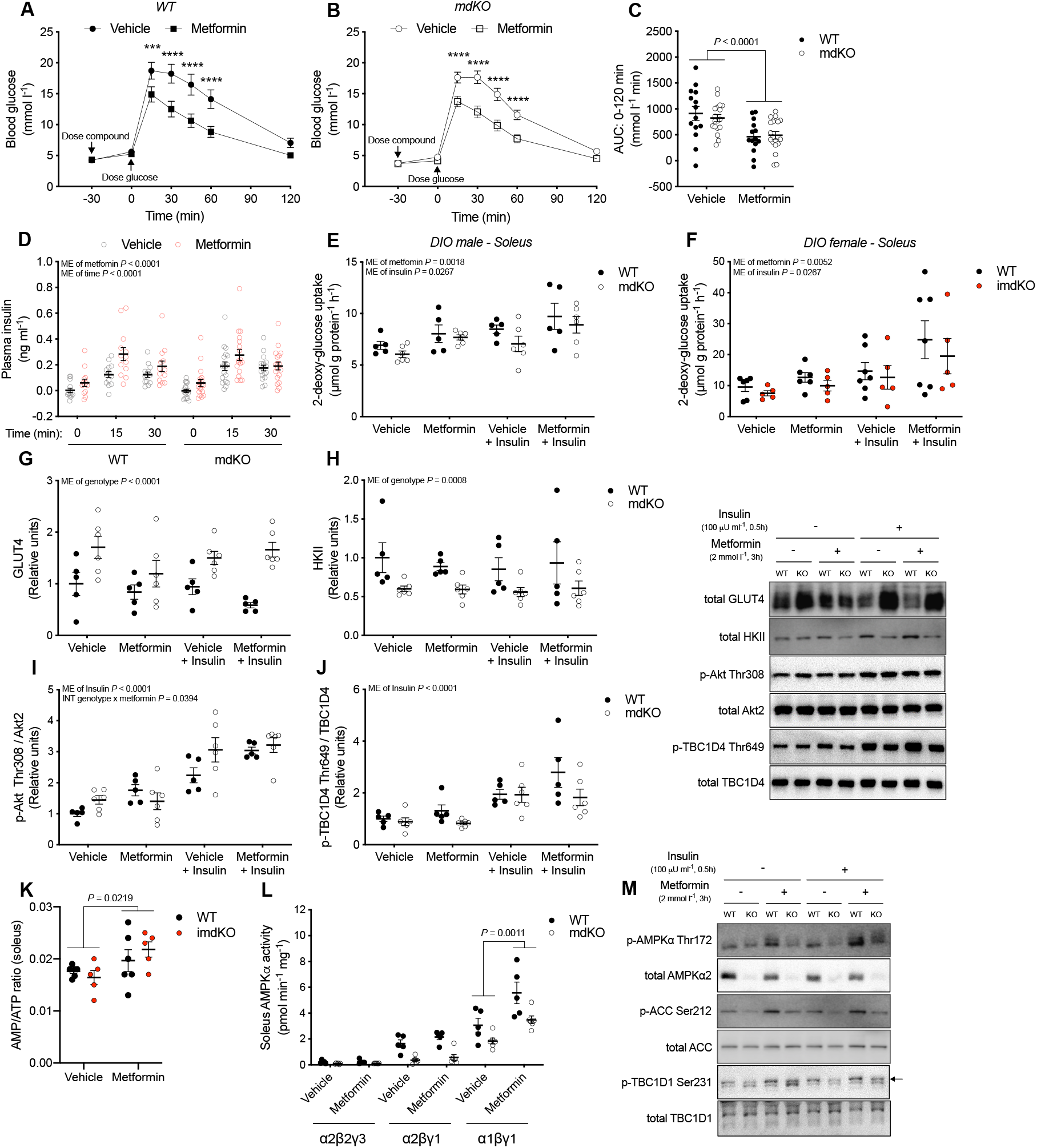
Acute metformin treatment improves whole body glucose tolerance and increases glucose uptake in isolated skeletal muscle from DIO mice independently of skeletal muscle AMPK. (A and B) Whole-body glucose tolerance test 30 min after i.p. dosed metformin (250 mg kg^-1^) in chow-fed male WT (A) and conventional muscle-specific AMPKα1α2 double KO (mdKO) mice (B). (C) Area under the curve (AUC) for the glucose tolerance test (0-120 min). (D) Plasma insulin levels at time points 0, 15, and 30 min during the glucose tolerance test. (E and F) Metformin- (2 mmol l^-1^) and submaximal insulin-stimulated (100 μU ml^-1^) 2-deoxyglucose uptake in isolated and incubated soleus muscle from DIO AMPK mdKO male mice (E), DIO imdKO female mice (F) and WT littermates. (G-J) Total protein expression of GLUT4 (G) and HKII (H), as well as phosphorylation of Akt Thr308 (I) and TBC1D4 Thr649 (J) in isolated and incubated soleus muscle from DIO AMPK mdKO male mice and WT littermates. Right: representative immunoblots. (K and L) AMP/ATP ratio (K) and AMPK activity of the α2βγ1, α1βγ1 and α2β2γ3 complexes (L) in isolated and incubated soleus muscle from DIO AMPK imdKO female mice (K), mdKO male mice (L) and WT littermates. (M) Representative immunoblots of AMPKα Thr172, ACC Ser212, and TBC1D1 Ser231 phosphorylation in isolated and incubated soleus muscle from male DIO muscle-specific AMPK mdKO mice and WT littermates. Data represent mean ± SEM. Data in (A-C, K and L) were analyzed using a two-way ANOVA while data in (D-J) were analyzed using a three-way ANOVA with Sídák post hoc analyses for multiple comparisons. *** denotes p<0.001 and **** denotes p<0.0001. ME, main effect. INT, interaction.

In an attempt to confirm previous observations in isolated rat skeletal muscle stimulated with a supra clinical concentration of metformin (Zhou et al., 2001), we isolated and incubated extensor digitorum longus (EDL) and soleus muscles from lean and DIO mice in the presence or absence of metformin (3h, 2 mmol l^-1^). Interestingly, acute metformin treatment increased glucose uptake in isolated soleus muscle from DIO mice independently of AMPK that was additive to the effect of a submaximal dose of insulin (Figures 3E, 3F and S3W-Z). These findings could not be explained by enhanced insulin signaling or increased protein expression of glucose transporter 4 (GLUT4) and hexokinase II (HKII) in metformin-treated WT and AMPK-deficient muscles (Figure 3G-J). Although the AMP/ATP ratio only increased in metformin-treated soleus muscles from DIO mice, AMPK activity and downstream signaling increased in all metformin-treated muscles (Figures 3K-M and S3AA-AG). Based on a recent study, this discrepancy may be explained by the PEN2-ATP6AP1 signaling axis that has been shown to activate AMPK in response to metformin without perturbing the AMP/ADP levels (Ma et al., 2022). The apparent discrepancy between unchanged glucose uptake/insulin sensitivity and increased AMPK activity in muscle from WT and AMPK-deficient mice likely stems from the predominant increase in AMPKα1βγ1-associated activity that corresponds to metformin-induced AMPK activation in non-muscle cells that are found in whole muscle tissue. Together, these findings demonstrate that prolonged treatment with a supra clinical concentration of metformin increases muscle glucose uptake by an AMPK-independent mechanism.

### Muscle AMPK is dispensable for the chronic actions of metformin

To investigate a possible role of muscle AMPK for mediating the effect of *in vivo* chronic metformin treatment, we administered metformin (p.o. 250 mg kg^-1^ day^-1^) once daily for a period of two weeks to WT and AMPK mdKO DIO mice and measured changes in blood glucose levels. The first and final acute dose of metformin decreased blood glucose levels similarly in WT and AMPK mdKO mice (Figures 4A, 4B, S4A, and S4B). However, the decrease in blood glucose levels was greater after the first dose than after the final dose likely due to a significant decrease in blood glucose levels after the treatment period (Figures 4C, 4D, S4C, and S4D). We did not observe that chronic metformin treatment affected food intake or body weight/composition compared to the vehicle-treated control mice (Figures S4E-H) as recently reported by others (Coll et al., 2020; Day et al., 2019). Glucose tolerance and *in vivo* insulin-stimulated glucose clearance were investigated ~24 hours after the final dose of metformin to examine potential glucoregulatory improvements by AMPK in response to chronic metformin treatment. At this time point metformin was still present in plasma and tissues (Figure 4E) although at ~50 to 250-fold lower levels compared to findings following acute treatment (Figure 2B). The decrease in blood glucose levels and increase in muscle and intestinal glucose clearance during insulin stimulation were not improved ~24 hours after the final dose of metformin (Figures 4F, 4G, and S4I) and AMPK activity/signaling in muscle, liver and intestine was not potentiated either (Figures 4H, 4I, and S4J-M). Intriguingly, glucose tolerance and plasma insulin levels seemed slightly compromised following chronic metformin treatment (Figures 4J-L and S4N and S4O), which indicates diverse effects of acute and chronic metformin treatment on glycemic control.

**Figure 4.**
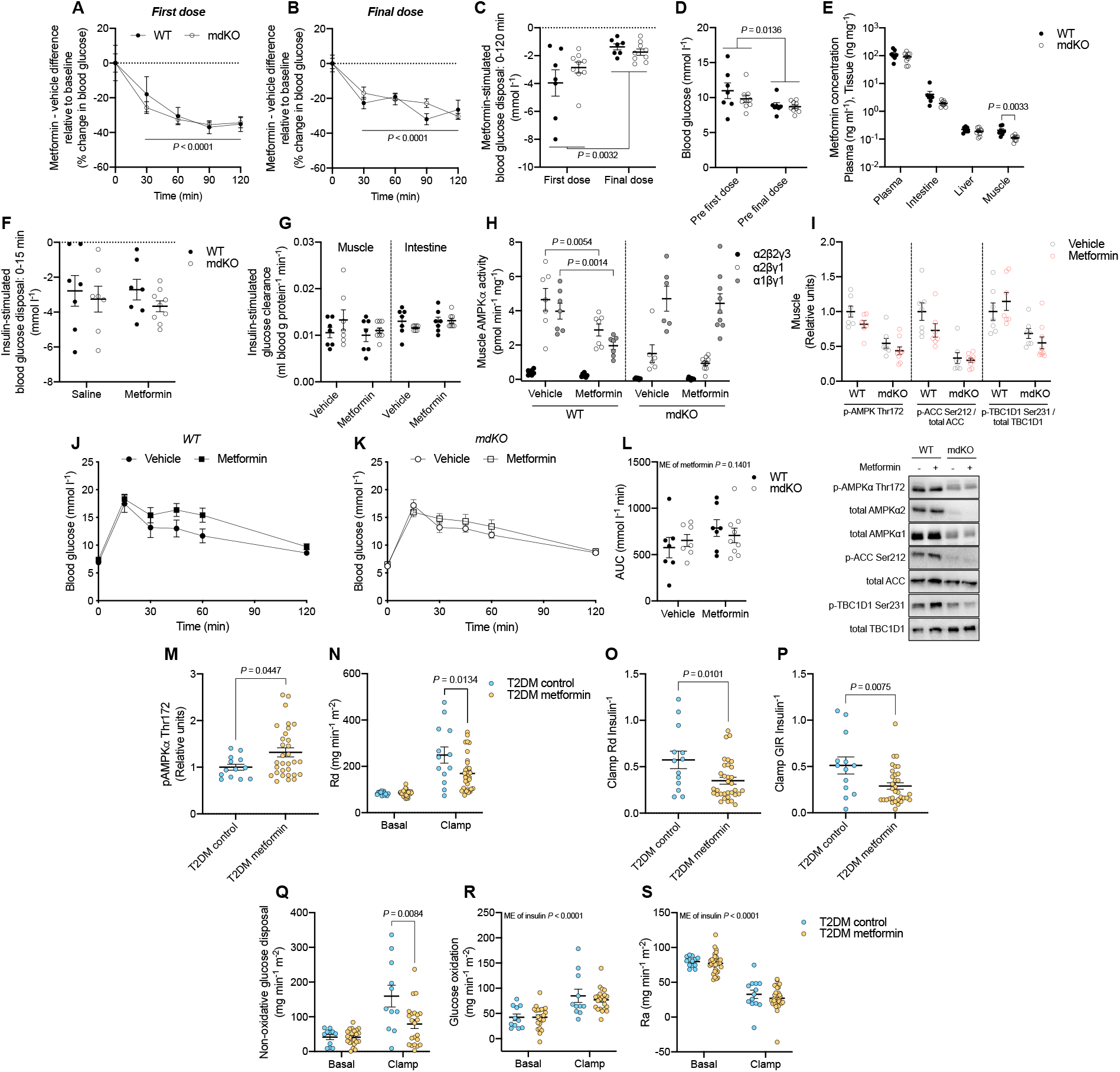
Elevated AMPK signaling in human skeletal muscle following chronic metformin treatment is not associated with enhanced peripheral insulin sensitivity. (A and B) Effect of orally dosed metformin (250 mg kg^-1^) vs. vehicle on blood glucose levels (%) in DIO AMPK mdKO mice and WT littermates. First dose (A) and final dose (B) of the chronic metformin treatment period (250 mg kg^-1^ day^-1^ for 14 days). (C) Acute metformin-stimulated blood glucose disposal (0-120 min) after the first and final dose of the chronic metformin treatment period. (D) Fed blood glucose levels immediately before the first and final dose of metformin of the treatment period. (E) Metformin concentration in plasma, intestine, skeletal muscle and liver tissue ~24 hours after the final dose of metformin. (F) Acute insulin-stimulated blood glucose disposal (0-15 min) ~24 hours after the final dose of vehicle/metformin. (G) Insulin-stimulated glucose clearance in quadriceps muscle and intestine from AMPK mdKO mice and WT littermates determined by the use of ^3^H-2-deoxy-D-glucose tracing (n = 7-9/7, mdKO/WT). (H-I) AMPK activity of the α2βγ1, α1βγ1 and α2β2γ3 complexes (H) as well as phosphorylation of AMPKα Thr172, ACC Ser212, and TBC1D1 Ser231 (I) in quadriceps muscle of chronic vehicle/metformin-treated DIO muscle-specific AMPKα1α2 KO mice and WT littermates. Below: representative immunoblots (J and K) Whole-body glucose tolerance ~24 hours after the final dose of vehicle/metformin in DIO WT (J) and AMPK mdKO mice (K). (L) Area under the curve (AUC) for the glucose tolerance test (0-120 min). (M) Phosphorylation of AMPKα Thr172 in muscle biopsy samples obtained from patients with type 2 diabetes mellitus treated with or without metformin. Metformin was withdrawn one week before the muscle biopsy sampling. (N-S) Measurements of Rd (N), Clamp Rd insulin^-1^ (O), Clamp GIR insulin^-1^ (P), Non-oxidative glucose disposal (Q), glucose oxidation (R), and Ra (S) during a hyperinsulinemic-euglycemic clamp in patients with type 2 diabetes mellitus (T2DM) treated with (yellow) or without (blue) metformin until one week before the clamp. Data represent mean ± SEM. Data in (E, I, and M-S) were analyzed using a homoscedastic twotailed student’s t-test. Data in (A-D, F-H, and J-K) were analyzed using a two-way ANOVA with (A-D, J and K) or without (F-H and L) repeated measures and Sídák post hoc analyses for multiple comparisons. Rd, glucose rate of disappearance. GIR, glucose infusion rate. Ra, glucose rate of appearance. ME, main effect.

### Elevated AMPK signaling in skeletal muscle from chronically metformin-treated type 2 diabetic patients is not associated with improved glycemic control

To provide human evidence for our findings in mice, we performed a retrospective analysis of four patient cohorts with T2DM. These had been treated with or without metformin up until one week before a hyperinsulinemic-euglycemic clamp (Højlund et al., 2009; Pedersen et al., 2015; Vind et al., 2011, 2012) (Table S1). We observed that phosphorylation of AMPKα Thr172 was increased in muscle biopsy samples obtained from the metformin-treated patients (Figures 4M and S4P). In contrast, the glucose infusion rate and the peripheral glucose disposal during insulin stimulation were decreased in the metformin-treated patients (Figures 4N-P and S4Q-S). This was driven by lower non-oxidative glucose disposal rates rather than a change in glucose oxidation or endogenous glucose production (Figures 4Q-S and S4T-V). Together, our findings indicate that elevated levels of AMPK activity in human skeletal muscle following chronic metformin treatment are not associated with enhanced peripheral insulin sensitivity.

## Discussion

Using lean and DIO mice with conventional and conditional KO of AMPKα1α2 in skeletal muscle, we report that *in vivo* activation of AMPK in mature skeletal muscle is not essential for the anti-hyperglycemic effect of metformin. Our findings of enhanced AMPKα Thr172 phosphorylation in skeletal muscle from chronically metformin-treated type 2 diabetic patients suggest that metformin has the potential to increase AMPK activity in human skeletal muscle. Yet, the lack of elevated basal glucose disposal rates in the metformin-treated type 2 diabetic cohorts together with observations of increased metformin-stimulated glucose uptake in muscle from AMPK-deficient mice certainly questions the role of muscle AMPK for the anti-hyperglycemic effect of metformin.

Pharmacologically-, contraction-, and exercise-induced activation of AMPK have been shown to enhance insulin-stimulated glucose uptake in skeletal muscle (Kjøbsted et al., 2015, 2017). Accordingly, metformin-induced activation of AMPK may drive improvements in muscle insulin sensitivity and evidence from several studies (Groop et al., 1989; Kim et al., 2002; Musi et al., 2002; Nosadini et al., 1987; Reaven et al., 1992; Riccio et al., 1991), but not all (Cusi et al., 1996; Natali et al., 2004; Wu et al., 1990), investigating whole-body glucose utilization during a hyperinsulinemic-euglycemic clamp in metformin-treated patients with T2DM supports such notion. However, only a few studies have reported effects of chronic metformin treatment on insulin-stimulated muscle glucose uptake in humans. These studies demonstrate that insulin-stimulated glucose uptake is not augmented, but rather tends to be diminished in skeletal muscle from overweight type 2 diabetic patients following metformin treatment (Hällsten et al., 2002; Karlsson et al., 2005). This is in line with our findings in mice and man demonstrating minor impairments in whole-body glucose tolerance and peripheral insulin-stimulated glucose disposal rates following chronic metformin treatment, respectively. Variation in dosing (McIntyre et al., 1991), patient characteristics (DeFronzo et al., 1991), metformin withdrawal (Diabetes Prevention Program Research Group, 2003), as well as time of last metformin ingestion prior to the experimental day (Færch et al., 2020; Musi et al., 2002) may explain observed discrepancies on glycemic control between studies.

Based on our findings demonstrating reduced liver glycogen and increased plasma glucagon levels in metformin-treated mice, it seems plausible that acute metformin treatment increases hepatic glucose production to counteract enhanced metformin-induced glucose disposal in other tissues as also recently proposed for non-diabetic individuals as well as individuals with recent-onset type 2 diabetes (Christensen et al., 2015; Gormsen et al., 2019; McCreight et al., 2020). Indeed, we found that the glucose clearance increased by ~75% in the intestine of acutely metformin-treated mice. This signifies that the intestine acts as a glucose sink following metformin treatment and thus plays a prominent role for the acute blood glucose-lowering effect of metformin (Bailey et al., 2008; Horakova et al., 2019; Rittig et al., 2021; Yang et al., 2021).

In conclusion, acute metformin treatment increases whole body glucose disposal, which occurs independently of AMPK-mediated glucose uptake in skeletal muscle but relies on enhanced intestinal glucose clearance. Furthermore, our observations in chronically metformin-treated patients with T2DM imply that metformin-induced activation of AMPK is insufficient to enhance basal and insulin-stimulated glucose uptake in human skeletal muscle.

## Acknowledgements

We thank all participants in the studies. The authors acknowledge the skilled technical help provided by Karina Olsen, Betina Bolmgren, and Irene Beck Nielsen (Department of Nutrition, Exercise, and Sports, Faculty of Science, University of Copenhagen). Funding for the study was provided by a research grant from the Danish Diabetes Academy, which is funded by the Novo Nordisk Foundation, grant number NNF17SA0031406 (to R.K.). Further funding for the study was provided by the EFSD/Lilly Young Investigator Research Award Programme (to R.K.) as well as by grants from the from the Danish Council for Independent Research (FSS: 8020-00288B), the Novo Nordisk Foundation (NNF21OC0070370) (to J.F.P.W.) and the Japan Society for the Promotion of Science, Grants-in-Aid for Scientific Research, grant number 19K20007 (to K.K.) and the EFSD/JDS Fellowship Program (to K.K.). None of the funding agencies had any role in the study design or in the collection or interpretation of the data.

## Author contributions

R.K. directed, managed and designed the animal work and performed the mouse experiments along with J.M.K., J.B.B., J.K.L., N.O.E. and K.K.. R.K. analyzed the animal and human data. R.K., J.B.B. and N.R.A. performed the biochemical analyses. M.F. and B.V. provided founder mice for the study. F.N., K.B. and Y.H. performed analyses of metformin and nucleotide levels. N.J. contributed with scientific expertise. K.H. and J.F.P.W. collected the human data. R.K. and J.F.P.W. wrote the manuscript. All authors interpreted the results, contributed to the discussion, edited and revised the manuscript, and finally read and approved the final version of the manuscript. R.K., K.H. and J.F.P.W are the guarantors of this work and, as such, had full access to all the data in the study and take responsibility for the integrity of the data and the accuracy of the data analysis.

## METHODS

### Experimental model and subject details

#### Ethical approvals

All animal experiments were approved by the Danish Animal Experiments Inspectorate (2013-15-2934-00911, 2014-15-2934-01037, and 2019-15-0201-01659) and complied with the EU convention for the protection of vertebrate animals used for scientific purposes (Council of Europe, Treaty 123/170, Strasbourg, France, 1985/1998). All human participants were given oral and written information after which written informed consent was obtained from all participants before entering the study. The study was approved by the relevant Local Ethics Committee of Denmark and was performed in accordance with the Helsinki Declaration.

#### Study participants

All participants were GAD65-antibody negative type 2 diabetic patients without signs of diabetic retinopathy, nephropathy, neuropathy, or macrovascular complications by blood test screening. Patients were treated by diet (n=10), diet and metformin (n=22), insulin (n=1), insulin and metformin (n=4), sulfonylurea (n=1), sulfonylurea and metformin (n=7), as well as sulfonylurea and rosiglitazone (n=1). Oral antidiabetics, antihypertensive and lipid lowering medication were withdrawn 1 week prior to the study and all participants were instructed to refrain from strenuous physical activity and exercise for a period of 48 hours before the hyperinsulinemic-euglycemic clamp. Data analyses related to this paper were performed post hoc. General patient information is provided in Table S1 while detailed patient and study information is available in previous publications (Højlund et al., 2009; Pedersen et al., 2015; Vind et al., 2011, 2012). Diabetes duration was not recorded in one study (Højlund et al., 2009) nor were measurements of glucose oxidation and non-oxidative glucose metabolism in a second study (Vind et al., 2011).

#### Animals

C57BL/6J male mice were purchased from Taconic Biosciences (Lille Skensved, Denmark). Female and male conventional AMPKα1α2 muscle-specific double KO (mdKO) mice were generated by mating double-floxed animals (AMPKα1^fl/fl^ and AMPKα2^fl/fl^) with double-floxed animals expressing the Cre-recombinase under the control of the human skeletal actin (HSA) promoter (AMPKα1^fl/fl^, AMPKα2^fl/fl^ and HSA-Cre^+/-^) (Fentz et al., 2015; Lantier et al., 2014). Double-floxed littermate animals were used as controls and referred to as WT mice. Female AMPKα1α2 inducible (conditional) muscle-specific double KO (imdKO) mice were generated by mating double-floxed animals (AMPKα1^fl/fl^ and AMPKα2^fl/fl^) with double floxed animals also expressing a tamoxifen-inducible Cre-recombinase under the control of the human skeletal actin (HSA) promoter (AMPKα1^fl/fl^, AMPKα2^fl/fl^ and HSA-MCM^+/-^) (Hingst et al., 2020). Acute deletion of AMPKα1 and AMPKα2 in skeletal muscle was achieved by i.p. injection of tamoxifen dissolved in 99% ethanol and resuspended in sunflower seed oil. Tamoxifen was provided to all mice (15-18 weeks of age) by a single injection (40 mg kg^-1^ body weight) on three occasions each separated by 48 h. This regimen produces an effective deletion of AMPKα1 and AMPKα2 in skeletal muscle three weeks after the final tamoxifen injection (Hingst et al., 2020). Double-floxed littermate animals were used as controls and referred to as WT mice. AMPK mdKO and imdKO mice and WT littermates were on a mixed background (C57BL/6J ~87.5% and SV129 ~12.5%). To confirm genotypes, genomic DNA from tail or ear snips was analyzed by PCR and muscle samples were analyzed for AMPKα1 and AMPKα2 protein expression by standard immunoblotting. All mice (except for wildtype C57BL/6J mice) were obtained from in-house breeding at a specific pathogen-free (SPF) animal facility. All mice were grouped-housed unless otherwise stated and acclimatized for at least 1 week at our local animal facility (12-h light/dark cycle (lights on 6 AM), 45-65% relative humidity, and 22 ±2°C) before use with access to water and standard rodent chow diet (Altromin no. 1324) *ad libitum*.

### Method details

#### Hyperinsulinemic euglycemic clamps

All participants underwent a hyperinsulinemic-euglycemic clamp following an overnight fast as described in detail previously (Højlund et al., 2009; Pedersen et al., 2015; Vind et al., 2011, 2012). In short, following a 2-h basal tracer equilibration, insulin was infused at a rate of 40 mU m^-2^ min^-1^ for 4 h or 80 mU m^-2^ min^-1^ for 3 h. A primed-constant [3-^3^H]-glucose infusion was used throughout the entire study and [3-^3^H]-glucose was added to the glucose infusates to maintain plasma specific activity constant at baseline as well as during the clamp. Plasma glucose concentrations were allowed to decline to ~5.5 mmol l^-1^ during the clamp before glucose infusion was initiated and hyperinsulinaemia was obtained at ~400 or ~900 pmol l^-1^ during the insulin infusion period. Whole body total glucose disposal rates (R_d_) and endogenous glucose production rates (R_a_) were calculated from the plasma [3-^3^H]-glucose specific activities using Steele’s non-steady-state equations as described (Hother-Nielsen et al., 1996). The distribution volume of glucose was set to 200 ml kg^-1^ body weight and the pool fraction was set to 0.65. The hyperinsulinemic-euglycemic clamps were combined with measurements of indirect calorimetry (TrueOne 2400 Metabolic Measurement System, Parvo Medics, USA) with the ventilated hood technique. Glucose oxidation rates were calculated using the stoichiometric equations of Frayn (Frayn, 1983), while non-oxidative glucose disposal rates were calculated by subtracting rates of glucose oxidation from R_d_.

#### Human muscle biopsies

Skeletal muscle biopsies were obtained from the m. Vastus Lateralis before and after the insulin infusion period using a modified Bergström needle with suction under local anesthesia (lidocaine). All muscle biopsies were obtained through separate incisions spaced 4-5 cm apart between the years 2007-2014. Biopsies were subsequently blotted and dissected free of blood, fat and connective tissue before frozen in liquid nitrogen. Lysates were prepared from freeze-dried (~48 h) muscle biopsies that were dissected free of visible fat, blood and connective tissue in a room with low humidity (25-30%).

#### High fat diets

Male C57Bl/6J mice were group-housed and placed on a standard 60% high fat diet (60% HFD; Research Diets no. D12492) for 16 weeks. All AMPK-deficient mice and WT littermates were single-caged on the day they were switched from standard rodent chow to a standard 60% high fat diet (60% HFD; Research Diets no. D12492). Following 6 weeks on the standard 60% HFD, mice were switched to a Western Diet containing 40% fat (40% WD, Research Diets no. D12079B) for an additional 6 weeks. Body weight was recorded once weekly.

#### Chronic metformin treatment of mice on HFD

Following 6 weeks on the standard 60% HFD and 4 weeks on the 40% WD, each mouse received a daily oral gavage of metformin dissolved in physiological saline (0.9%) or vehicle for 14 days in the early part of the light cycle (~7 AM to 10 AM). Food intake (40% WD) and body weight were recorded daily. Blood glucose levels were determined from the tail vein at *t* = 0, 30, 60, 90 and 120 min using a blood glucometer (Contour XT, Bayer) on the 1^st^ and 14^th^ day of treatment. In the morning on day 15, approximately 24 hours after the 14^th^ dose of metformin, all mice underwent an oral glucose tolerance test (for details see below). On day 17 and 18, approximately 24 hours after the 16^th^ and 17^th^ dose of metformin, in vivo insulin-stimulated glucose clearance was determined in half of the animal study cohort (for details see below).

#### Body composition

Body composition was measured in conscious mice using magnetic resonance imaging (EchoMRI-4in1TM, Echo Medical System).

#### Metformin tolerance tests

Metformin dissolved in physiological saline (0.9%) was administered to single-housed and nonfasted animals in the early part of their light cycle by a single intraperitoneal injection or by oral gavage (250 mg kg^-1^). Blood glucose levels were determined from the tail vein at *t* = 0, 30, 60 and 75 min using a blood glucometer (Contour XT, Bayer). At *t* = 45, animals were anaesthetized by a single intraperitoneal injection of pentobarbital and xylocain (10 and 0.5 mg per 100 g of body weight, respectively) dissolved in physiological saline (0.9%). At *t* = 60 or *t* = 75, the thorax cavity was exposed and blood was drawn by cardiac puncture using heparinized syringes and collected into standard as well as EDTA-coated tubes after which tissues were dissected, frozen in liquid nitrogen and stored at −80°C until further analyses. All liver tissue samples were freeze-clamped in situ using liquid nitrogen-cooled flat tongs to avoid liver metabolic disturbances following dissection.

#### In vivo metformin-stimulated glucose clearance

Metformin-stimulated glucose clearance was evaluated during metformin tolerance tests lasting 75 min. At *t* = 60 min, a bolus of [^3^H]2-deoxy-glucose (12.3 MBq kg^-1^ of body weight) dissolved in physiological saline (0.9%) was administrated to anaesthetized animals by a single retro-orbital injection. To determine glucose clearance during metformin stimulation, blood glucose levels were determined from the tail vein at *t* = 60, 65, 70, and 75 min using a glucometer (Contour XT, Bayer). Moreover, 25 μl of blood was collected at *t* = 60, 65, 70, and 75 min and transferred to separate tubes containing 60 μl of BaOH. Blood-BaOH mixture samples were immediately vortexed after which 60 μl of ZnSO_4_ was added followed by a second vortex. Following the experiment, all blood mixture samples were centrifuged at 14,000 *g* for 5 min at room temperature. Radioactivity of [^3^H]2-deoxy-glucose was measured in 50 μl of supernatant from the blood mixture samples by liquid scintillation counting and used to assess tissue glucose clearance.

#### Oral glucose tolerance test after chronic metformin treatment

On the day of the last dose of metformin (day 14), animals were single housed and fasted overnight for 16 h (~3 PM to 7 AM) before they were administered with an oral gavage of glucose (2 g/kg body weight) dissolved in physiological saline (0.9%). Blood glucose was determined from the tail vein at *t* = 0, 15, 30, 45, 60, and 120 min using a blood glucometer (Contour XT, Bayer). At *t* = 0, 15, and 30 min, ~35 μl of blood was drawn into heparinized capillary tubes to determine plasma insulin concentrations.

#### In vivo insulin-stimulated glucose clearance

Non-fasted animals were anaesthetized by a single intraperitoneal injection of pentobarbital and xylocain (9 and 0.5 mg 100 g^-1^ of body weight, respectively) dissolved in physiological saline (0.9%) and left to recover on a heating plate (30°C) for ~15 min. Subsequently, a bolus of [^3^H]2-deoxyglucose (12.3 MBq kg^-1^ of body weight) and insulin (1 Unit kg^-1^ of body weight) dissolved in physiological saline (0.9%) was administrated by a single retro-orbital injection. To determine glucose clearance during insulin stimulation, blood glucose levels were determined from the tail vein at *t* = 0, 5, 10 and 15 min using a blood glucometer (Contour XT, Bayer). Moreover, 25 μl of blood was collected at *t* = 0, 5, 10 and 15 min and transferred to separate tubes containing 60 μl of BaOH. Blood-BaOH mixture samples were immediately vortexed after which 60 μl of ZnSO_4_ was added followed by a second vortex. At *t* = 15 the thorax cavity was exposed and blood was drawn by cardiac puncture using heparinized syringes and collected into standard as well as EDTA-coated tubes after which tissues were dissected, frozen in liquid nitrogen and stored at −80°C until further analyses. All liver tissue samples were freeze-clamped in situ using liquid nitrogen-cooled flat tongs to avoid liver metabolic disturbances following dissection. After the experiment, all blood mixture samples were centrifuged at 14,000 *g* for 5 min at room temperature. Radioactivity of [^3^H]2-deoxy-glucose was measured in 50 μl of supernatant from all the blood mixture samples by liquid scintillation counting and used to assess tissue glucose clearance.

#### Combined metformin and glucose tolerance test

Animals were single housed and fasted overnight for 16 h (~3 PM to ~7 AM) after which metformin-HCl (250 mg kg^-1^) dissolved in physiological saline (0.9%) was administered by a single intraperitoneal injection or by oral gavage (*t* = −30 min). Before and 30 min after metformin administration (*t* = −30 and 0 min), blood glucose levels were determined and glucose (2 g/kg body weight) dissolved in physiological saline (0.9%) was administered by a single intraperitoneal injection. Blood glucose levels were determined at 15, 30, 45, 60, and 120 min. At *t* = 0, 15, and 30 min, ~35 μl of blood was drawn into heparinized capillary tubes to determine plasma insulin concentrations.

#### 2-deoxyglucose uptake in isolated skeletal muscle

Non-fasted animals fed either chow or HFD were anaesthetized by i.p. injection of pentobarbital and xylocain (10 and 0.5 mg 100 g^-1^ of body weight, respectively) dissolved in physiological saline (0.9%). Next, soleus and extensor digitorum longus (EDL) muscles were dissected and suspended at resting tension in incubation chambers (model 610/820M; Danish Myo Technology, Denmark) containing Krebs-Ringer-Henseleit (KRH) buffer (117 mmol l^-1^ NaCl, 4.7 mmol l^-1^ KCl, 2.5 mmol l^-1^ CaCl_2_, 1.2 mmol l^-1^ KH_2_PO_4_, 1.2 mmol l^-1^ MgSO_4_, 0.5 mmol l^-1^NaHCO_3_, pH 7.4) supplemented with 0.1% BSA, 8 mmol l^-1^ mannitol, and 2 mmol l^-1^ pyruvate. The incubation buffer was continuously oxygenated (95% O_2_ and 5% CO_2_) and maintained at 30°C throughout the incubation period. After 10 min pre-incubation, all muscles were incubated for 3 h in KRH buffer with or without metformin (2 mmol l^-1^). During this period, the KRH incubation buffer was replaced every hour. Subsequently, vehicle and metformin-stimulated 2-deoxyglucose uptake was measured during the last 10 min of the 3 h stimulation period by adding 1 mmol l^-1^ [^3^H]2-deoxyglucose (0.028 MBq ml^-1^), 7 mmol l^-1^ [^14^C]mannitol (0.083 MBq ml^-1^), and 2 mmol l^-1^ pyruvate to the KRH buffer with or without metformin (2 mmol l^-1^). For measurements of insulin-stimulated 2-deoxyglucose uptake, muscles were incubated for 30 min in KRH buffer in the absence or presence of a submaximal dose (100 μU ml^-1^) of insulin (Actrapid, Novo Nordisk, Denmark) following the 3 h stimulation period. Insulin-stimulated 2-deoxyglucose uptake was measured during the last 10 min of the 30 min stimulation period by adding 1 mmol l^-1^ [^3^H]2-deoxyglucose (0.028 MBq ml^-1^), 7 mmol l^-1^ [^14^C]mannitol (0.083 MBq ml^-1^) (Hartmann Analytic), and 2 mmol l^-1^ pyruvate to the KRH buffer containing no insulin or a submaximal insulin concentration (100 μU ml^-1^). After the incubation period, muscles were washed in ice-cold KRH buffer, dried on filter paper and frozen in liquid nitrogen. Muscle 2-deoxyglucose uptake was assessed by the accumulation of [^3^H]2-deoxyglucose in muscle using [^14^C]mannitol as an extracellular marker. Radioactivity was measured on 150 or 250 μl of muscle lysate by liquid scintillation counting (Ultima Gold and Tri-Carb 4910 TR, Perkin Elmer) and related to the specific activity of the incubation buffer.

#### Metformin concentration

Metformin concentration in plasma and tissue samples was analyzed at the Department of Clinical Pharmacology, Pharmacy and Environmental Medicine, Institute of Public Health, University of Southern Denmark, by the use of liquid chromatography and triple quadrupole mass spectrometry (LC–MS/MS) according to previously published methods (Kuhlmann et al., 2021; McCreight et al., 2018). The intra- and inter-day variability was <8%. The limit of detection (LOD) for the method was 1 ng ml^-1^ and limit of quantification was 10 ng ml^-1^. The tissue samples were homogenized during the extraction procedure by the following method: 30 mg of tissue was accurately weighed into a tarred 2 ml polypropylene cryo-protected tube (SARSTEDT), and 100 μl Milli-Q treated water, 10 μl 25 μg ml^-1^ metformin-(dimethyl-d6) hydrochloride (internal standard, Sigma-Aldrich), 20 μl 0.53 mol l^-1^ ammonium acetate and 390 μl acetonitrile were added. Three stainless steel beads (3 mm) were added into the microtubes and the samples were homogenized for 2 x 3 minutes at 25 Hz using a Mixer Mill MM 400 (Retsch). The samples were hereafter centrifuged at 21,000 g for 20 minutes. A volume of 10 μL of the supernatant was injected into the LC-MS/MS system. The concentration of metformin in the tissue samples was reported as ng g^-1^ tissue.

#### Tissue metabolites (nucleotides and lactate)

Approximately 5-10 mg of wet weight mouse tissue (skeletal muscle, liver and small intestine) was extracted with 1 mmol l^-1^ perchloric acid, neutralized with 1 mmol l^-1^ Potassium Hydroxide and 0.4 mmol l^-1^ Tris-base and subsequently analyzed for purine nucleotides by reverse-phase high-performance liquid chromatography as previously described (Tullson et al., 1990). Lactate concentration in extracted and neutralized tissues was determined by a fluorometric method (Lowry and Passonneau, 1972) and given as mmol kg^-1^ wet weight.

#### Plasma metabolites

Plasma was prepared from whole blood by centrifugation at 16,000 *g* for 5 min at 4°C and stored at −80°C until further analyses. Plasma lactate was assayed using a lactate meter (Lactate Plus, Novo Biomedical, UK). Plasma insulin and glucagon were determined using an ELISA from ALPCO and Mercodia, respectively.

#### Tissue homogenization

Tissues were pulverized in liquid nitrogen prior to homogenization. Tissues were homogenized in ice-cold lysis buffer (10% glycerol, 20 mmol l^-1^ sodium pyrophosphate, 1% NP-40, 2 mmol l^-1^ phenylmethylsulfonyl fluoride [PMSF], 150 mmol l^-1^ sodium chloride, 50 mmol l^-1^ HEPES, 20 mmol l^-1^ b-glycerophosphate, 10 mmol l^-1^ sodium fluoride, 1 mmol l^-1^ EDTA, 1 mmol l^-1^ EGTA, 10 mg ml^-1^ aprotinin, 3 mmol l^-1^ benzamidine, 10 mg ml^-1^ leupeptin, and 2 mmol l^-1^ sodium orthovanadate, pH 7.5) using steel beads and a TissueLyzer II (QIAGEN). Tissue homogenates were rotated end-over-end for 1 h at 4°C before part of the homogenates were centrifuged at 16,000 *g* for 20 min at 4°C to generate tissue lysates (supernatant). Tissue homogenates and lysates were collected, frozen in liquid nitrogen and stored at −80°C until further analyses. Total protein abundance in tissue homogenates and lysates were determined in triplicates by the bicinchoninic acid method (Thermo Fisher Scientific).

#### Immunoblotting

Tissue lysate samples were adjusted to equal protein concentrations using Milli-Q water and denatured by mixing with 6x Laemmli sample buffer and heating at 96°C for 10 min. Equal amounts of protein were separated by SDS-PAGE on self-cast gels and subsequently transferred to polyvinylidene fluoride (PVDF) membranes using semi-dry blotting. After blocking the membranes for 10 min in TBS-T containing 2% non-fat milk, membranes were incubated in primary antibody over night at 4°C. The following day, membranes were rinsed in TBS-T for 5 min and then incubated 1 h with HRP-conjugated secondary antibody at room temperature. Following a second rinse in TBS-T (~30 min), membrane-bound proteins were visualized with chemiluminescence using a digital imaging system (ChemiDoc MP System, BioRad) and quantified using Image Lab (BioRad). Protein Plus Precision All Blue standards were used as marker of molecular weight (BioRad) and a tissue lysate standard were loaded twice on each gel to control for transfer efficiency. Linearity was assessed for all measured proteins to confirm that the band intensities of each membrane were within the dynamic range.

#### AMPK activity assay

AMPKα2β2γ3-associated activity was measured on AMPKγ3 immunoprecipitates (IPs) that were prepared from equal amounts of tissue protein lysate, AMPKγ3-specific antibody, Protein G Agarose beads (Millipore) and AMPK-IP buffer (50 mmol l^-1^ NaCl, 1% Triton X-100, 50 mmol l^-1^ sodium fluoride, 5 mmol l^-1^ sodium-pyrophosphate, 20 mmol l^-1^ Tris-base, pH 7.5, 500 mmol l^-1^ PMSF, 2 mmol l^-1^ dithiothreitol, 4 mg ml^-1^ leupeptin, 50 mg ml^-1^ soybean trypsin inhibitor, 6 mmol l^-1^ benzamidine, and 250 mmol l^-1^ sucrose). Following end-over-end rotation over night at 4°C, IPs were centrifuged at 500 *g* for 1 min and subsequently washed once in AMPK-IP buffer, once in 6x assay buffer (240 mmol l^-1^ HEPES, 480 mmol l^-1^ NaCl, pH 7.0), and twice in 3x assay buffer. The AMPK activity assay ran for 30 min at 30°C in 30 μl kinase mix (40 mmol l^-1^ HEPES, 80 mmol l^-1^ NaCl, 833 μmol l^-1^ dithiothreitol, 200 μmol l^-1^ AMP, 100 μmol l^-1^ AMARA peptide (Schafer-N), 5 mmol l^-1^ MgCl_2_, 200 μmol l^-1^ ATP, and 2 μCi of [γ-33P]-ATP; Hartmann Analytic). The AMPK kinase reaction was terminated by the adding 10 μl of 1% phosphoric acid to each sample. Twenty microliters of the reaction mix were subsequently spotted on P81 filter paper (Saint Vincent’s Institute, Medical Research, AUS). Next, filter papers were washed thrice in 1% phosphoric acid for 15 min. Radioactivity was analyzed on dried filter paper using a Typhoon FLA 7000 IP (GE Healthcare). AMPKα2βγ1-associated activity was measured on supernatant from the AMPKγ3 IPs using a AMPKα2-specific antibody for a second IP while AMPKα1βγ1-associated activity was measured on supernatants from the AMPKα2 IPs using a AMPKα1-specific antibody for a third IP. In liver and intestinal tissues, AMPK activity was only assessed by AMPKα2 and AMPKα1 IPs since AMPKγ3 protein is not expressed in these tissues.

#### Tissue glycogen content

Equal amounts of muscle (400 μg) and liver (100 μg) protein homogenate were added to separate tubes containing 2 mmol l^-1^ hydrochloride and heated at 96°C for 2 h. Tissue glycogen content was determined by a fluorometric method (Lowry and Passonneau, 1972) and given as glycosyl units per total amount of protein.

#### Calculation of metformin- and insulin-stimulated tissue glucose clearance

Equal amounts of tissue homogenate were transferred to a set of tubes containing 4.6% perchloric acid (1:13 v/v) as well as a set of other tubes containing 0.3 M BaOH (1:6.5 v/v). Both set of tubes were then vortexed after which 0.3 M ZnSO_4_ was added to the tubes containing BaOH (1:6.5:6.5 v/v). Following vortexing, both set of tubes were centrifuged at 13,000 *g* for 4 min at room temperature and [^3^H] radioactivity of the supernatant was determined by liquid scintillation counting (Ultima Gold and Tri-Carb 2910 TR, Perkin Elmer). The amount of phosphorylated [^3^H]2-deoxy-glucose ([^3^H]2-deoxy-glucose-6-P) cleared by the tissues was calculated by subtracting the [^3^H] radioactivity of the BaOH/ZnSO_4_ supernatant fraction (contains only [^3^H]2-deoxy-glucose) from that of the perchloric acid supernatant fraction (contains [^3^H]2-deoxy-glucose and [^3^H]2-deoxy-glucose-6-P). To calculate the rate of tissue glucose clearance, the radioactivity of [^3^H]2-deoxy-glucose-6-P was related to the amount of total protein in the samples, the specific activity of the blood, the time of exposure and the average blood glucose concentration of each individual animal. Glucose clearance was given as ml blood^-1^ mg protein^-1^ min^-1^.

### Quantification and statistical analysis

An unpaired two-tailed Student’s t test was used for comparisons between two groups. Two and three independent variables were compared using a two-way and a three-way ANOVA with or without repeated measures, respectively, followed by Sidák’s multiple comparisons test. The p-value < 0.05 was considered to be statistically significant. Data are presented as mean ± SEM. No methods were used to determine whether the data met assumptions of the statistical approach. Statistical parameters can be found in the individual figure legends. Statistical analyzes were performed using GraphPad Prism 9.0 (GraphPad Software).

