## Supplementary material for "Metformin improves glycemia independently of skeletal muscle AMPK via enhanced intestinal glucose clearance": Table S1

**Table S1. Characteristics of the patients.**

| Variable | T2DM Control (n=13) | T2DM Metformin (n=33) | P-value |
|---|---|---|---|
| Age - yr | 53.4 ± 2.0 | 53.1 ± 0.9 | 0.881 |
| Men - no (%) | 11 (84.6) | 26 (78.8) | 0.662 |
| Weight - kg | 96.6 ± 4.7 | 98.5 ± 2.5 | 0.703 |
| Body mass index | 31.8 ± 0.7 | 30.9 ± 1.3 | 0.507 |
| HbA1c - % | 7.0 ± 0.4 | 7.2 ± 0.2 | 0.587 |
| Plasma glucose, basal - mmol $l^{-1}$ | 8.2 ± 0.5 | 9.5 ± 0.4 | 0.056 |
| Plasma glucose, clamp - mmol $l^{-1}$ | 5.3 ± 0.1 | 5.3 ± 0.1 | 0.862 |
| Serum insulin, basal - pmol $l^{-1}$ | 59.9 ± 9.6 | 69.6 ± 6.7 | 0.435 |
| Serum insulin, clamp - pmol $l^{-1}$ | 477.2 ± 55.5 | 571.6 ± 44.3 | 0.238 |
| Diabetes duration - yr | 4.3 ± 1.5 | 4.3 ± 0.7 | 0.990 |
| *Exclusion of patients receiving anti-diabetic medication other than metformin (Control, n=3; Metformin, n=11)* | | | |
| Age - yr | 54.7 ± 2.4 | 52.8 ± 1.0 | 0.416 |
| Men - no (%) | 9 (90.0) | 18 (81.8) | 0.569 |
| Weight - kg | 100.6 ± 5.1 | 97.4 ± 3.5 | 0.605 |
| Body mass index | 31.7 ± 1.5 | 31.4 ± 0.9 | 0.894 |
| HbA1c - % | 6.9 ± 0.4 | 7.1 ± 0.2 | 0.598 |
| Plasma glucose, basal - mmol $l^{-1}$ | 7.7 ± 0.5 | 9.1 ± 0.4 | 0.033 |
| Plasma glucose, clamp - mmol $l^{-1}$ | 5.2 ± 0.1 | 5.3 ± 0.1 | 0.626 |
| Serum insulin, basal - pmol $l^{-1}$ | 61.2 ± 11.3 | 64.5 ± 7.8 | 0.816 |
| Serum insulin, clamp - pmol $l^{-1}$ | 475.6 ± 71.1 | 576.1 ± 53.7 | 0.289 |
| Diabetes duration - yr | 2.8 ± 0.8 | 3.1 ± 0.6 | 0.729 |

Values represent means ± SEM. Data were analysed by a homoscedastic two-tailed student's t-test. The upper pat of the table represents data from patients treated with or without metformin and other anti-diabetic medication (e.g. insulin). The lower part of the table represents data from patients only treated with or without metformin.

### Supplemental figures, figure titles and legends

#### Figure S1

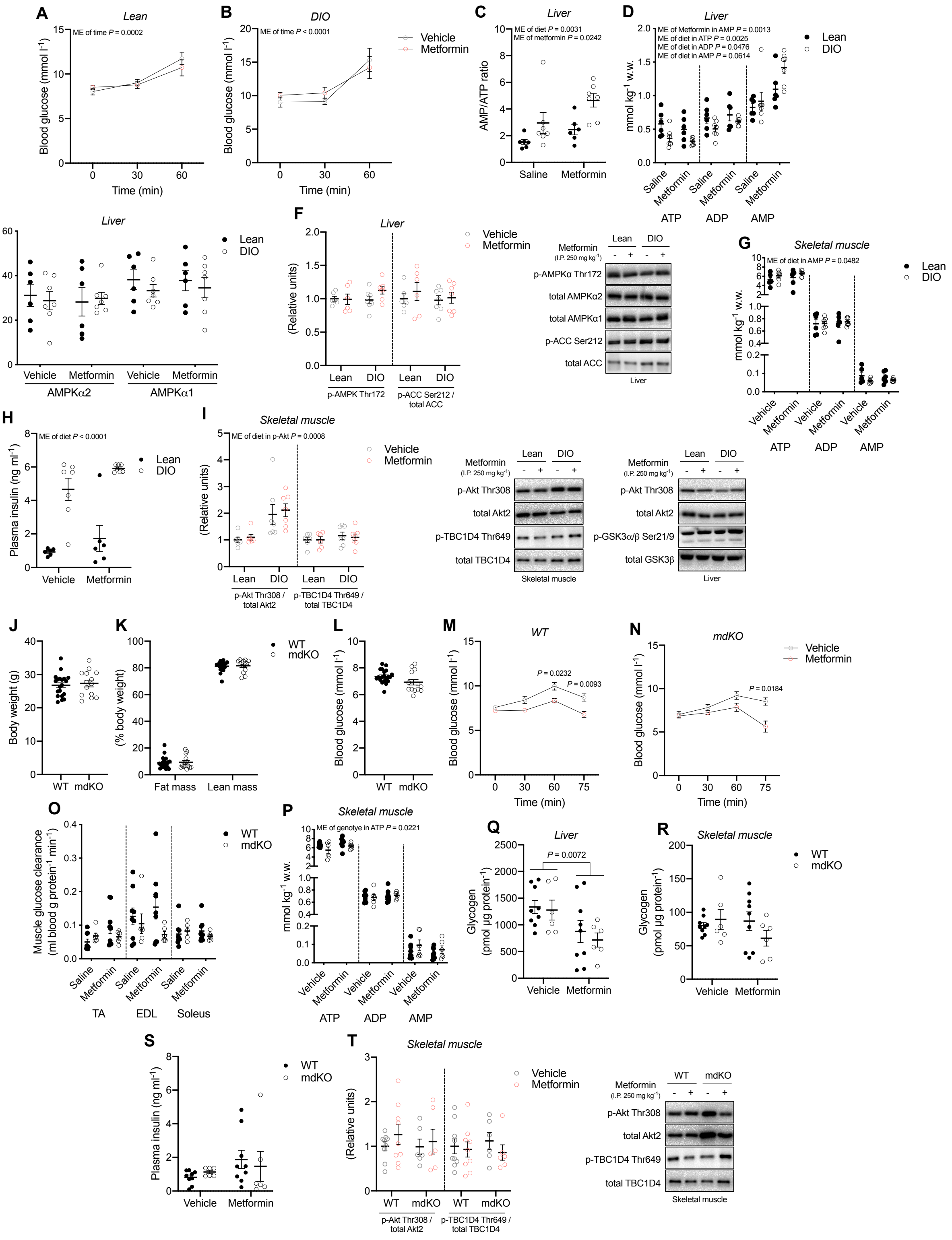

**Figure S1. Acute metformin treatment lowers blood glucose levels in lean and DIO mice independently of enhanced AMPK activity in skeletal muscle, Related to Figure 1.**

(A and B) Effect of i.p. administered metformin (250 mg $kg^{-1}$) vs. vehicle on blood glucose levels (mmol $l^{-1}$) in lean and DIO mice (n = 6/7, chow/HFD).

(C-F) AMP/ATP ratio (C) ATP, ADP and AMP levels (D), AMPK activity of the α2- and α1-associated complexes (E) as well as phosphorylation of AMPKα Thr172 and ACC Ser212 (F) in liver of vehicle- and metformin-treated lean and DIO mice. Right: representative immunoblots.

(G) ATP, ADP and AMP levels in quadriceps muscle of vehicle- and metformin-treated mice.

(H) Plasma insulin levels 60 min after metformin treatment of lean and DIO mice.

(I) Phosphorylation of Akt Thr308 and TBC1D4 Thr649 in quadriceps muscle (quantified) and liver of vehicle- and metformin-treated lean and DIO mice. Right: representative immunoblots.

(J-L) Body weight (J), body composition (K) and fed blood glucose (L) in lean conventional muscle-specific AMPKα1α2 KO (mdKO) female mice and wildtype (WT) littermates measured in the morning after a night of free access to chow diet (n = 20/14, WT/mdKO).

(M and N) Effect of i.p. administered metformin (250 mg $kg^{-1}$) vs. vehicle on blood glucose levels (mmol $l^{-1}$) in WT and mdKO chow-fed mice (n = 6-11 per group).

(O) Glucose clearance in tibialis anterior (TA), extensor digitorum longus (EDL) and soleus muscle of vehicle- and metformin-treated AMPK mdKO mice and WT littermates determined by the use of $^{3}H$-2-deoxy-D-glucose tracing (n = 5-9 per group).

(P) ATP, ADP and AMP levels in quadriceps muscle of vehicle- and metformin-treated WT and mdKO mice.

(Q and R) Glycogen levels (glycosyl units) in liver (Q) and quadriceps muscle (R) of vehicle- and metformin-treated WT and mdKO mice.

(S) Plasma insulin levels 75 min after metformin treatment in WT and mdKO mice.

(T) Phosphorylation of Akt Thr308 and TBC1D4 Thr649 in quadriceps muscle of vehicle- and metformin-treated WT and mdKO mice. Right: representative immunoblots.

Data represent mean ± SEM. Data in panels J, K, and L were analyzed using a homoscedastic two-tailed student's t-test. Data in the remaining panels were analyzed using a two-way ANOVA with (A, B, M and N) or without (C, D, E, F, G, H, I, O, P, Q, R, S, and T) repeated measures and Sídák post hoc analyses for multiple comparisons. AU, arbitrary units. w.w, wet weight. ME, main effect.

**Figure S2**

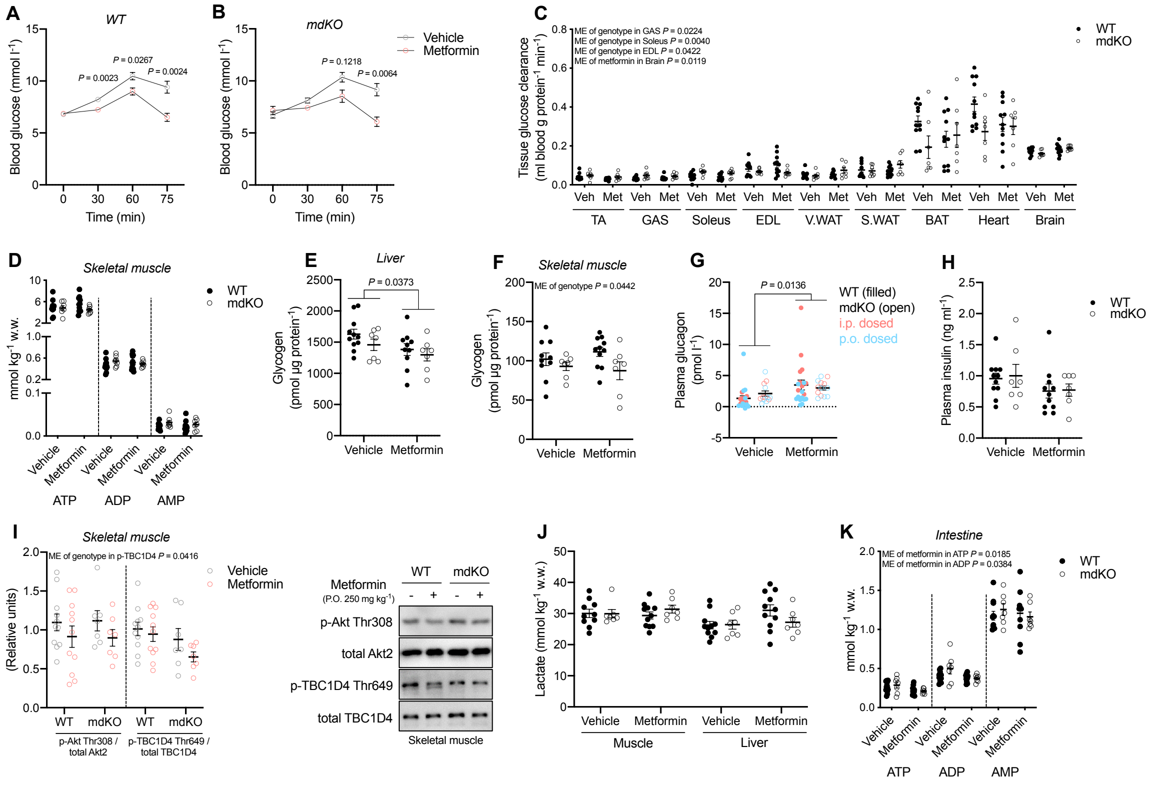


**Figure S2. Acute metformin treatment lowers blood glucose levels by enhancing intestinal glucose clearance, Related to figure 2.**

(A and B) Effect of p.o. administered metformin (250 mg $kg^{-1}$) vs. vehicle on blood glucose levels (mmol $l^{-1}$) WT and mdKO chow-fed mice (n = 11/7, WT/mdKO).

(C) Glucose clearance in tibialis anterior muscle (TA), gastrocnemius muscle (GAS), soleus muscle, extensor digitorum longus muscle (EDL), visceral white adipose tissue (V.WAT), subcutaneous white adipose tissue (S.WAT), brown adipose tissue (BAT), heart and brain tissue of orally vehicle- and metformin-treated AMPK mdKO mice and WT littermates determined by the use of $^{3}$H-2-deoxy-D-glucose tracing (n = 11/7, WT/mdKO).

(D) ATP, ADP and AMP levels in quadriceps muscle of orally vehicle- and metformin-treated WT and mdKO mice.

(E and F) Glycogen levels (glycosyl units) in liver (E) and quadriceps muscle (F) of orally vehicle- and metformin-treated WT and mdKO mice.

(G) Plasma glucagon levels of i.p. (red colour) and p.o. (blue colour) vehicle- and metformin-treated WT (filled dots) and mdKO (open dots) mice 75 min after treatment.

(H) Plasma insulin levels 75 min after orally metformin treatment of WT and mdKO mice.

(I) Phosphorylation of Akt Thr308 and TBC1D4 Thr649 in quadriceps muscle of orally vehicle- and metformin-treated WT and mdKO mice. Right: representative immunoblots.

(J and K) Lactate levels in quadriceps muscle and liver tissue (J) as well as ATP, ADP and AMP levels in the intestine (K) of orally vehicle- and metformin-treated WT and mdKO mice. Data represent mean ± SEM. Data in all panels were analyzed using a two-way ANOVA with (A and B) or without (C-K) repeated measures and Sídák post hoc analyses for multiple comparisons. w.w, wet weight. ME, main effect.

**Figure S3**

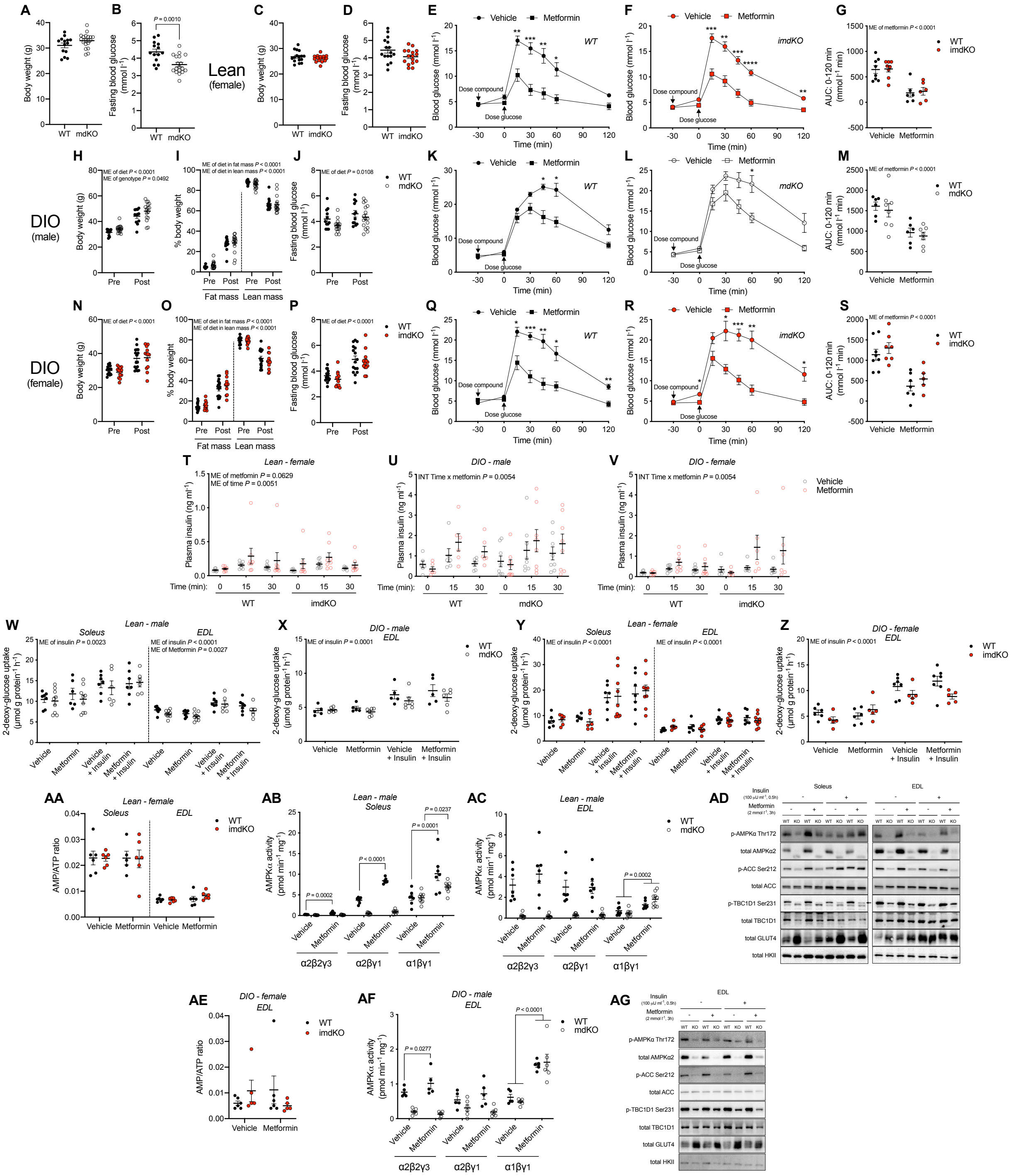


**Figure S3. Acute metformin treatment improves whole body glucose tolerance and increases glucose uptake in isolated skeletal muscle from DIO mice independently of skeletal muscle AMPK, Related to figure 3.**

(A and B) Body weight (A) and fasting blood glucose (B) in lean chow-fed WT and mdKO male mice measured in the morning after overnight fasting (n = 14/18, WT/mdKO).

(C and D) Body weight (A) and fasting blood glucose (B) in chow-fed WT and inducible (conditional) muscle-specific AMPKα1α2 double KO (imdKO) female mice measured in the morning after overnight fasting (n = 16 in both groups).

(E and F) Whole-body glucose tolerance 30 min after i.p. dosed metformin (250 mg $kg^{-1}$) in lean chow-fed WT (E) and imdKO (F) female mice.

(G) Area under the curve (AUC) for the glucose tolerance tests in (E) and (F) (0-120 min).

(H-J) Body weight (H), body composition (I) and fasting blood glucose (J) measured in mdKO male mice and WT littermates before (Pre) and after (Post) the HFD intervention in the morning after overnight fasting (n = 12/16, WT/mdKO).

(K and L) Whole-body glucose tolerance 30 min after p.o. dosed metformin (250 mg $kg^{-1}$) in DIO WT (K) and mdKO (L) male mice.

(M) Area under the curve (AUC) for the glucose tolerance tests in (K) and (L) (0-120 min).

(N-P) Body weight (N), body composition (O) and fasting blood glucose (P) measured in imdKO female mice and WT littermates before (Pre) and after (Post) the HFD intervention in the morning after overnight fasting (n = 16/13, WT/imdKO).

(Q and R) Whole-body glucose tolerance 30 min after i.p. dosed metformin (250 mg $kg^{-1}$) in DIO WT (Q) and imdKO (R) female mice.

(S) Area under the curve (AUC) for the glucose tolerance tests in (Q) and (R) (0-120 min).

(T-V) Plasma insulin levels at time points 0, 15, and 30 min during the glucose tolerance test in panels E and F (T), K and L (U) as well as Q and R (V).

(W-Z) Metformin- (2 mmol $l^{-1}$) and submaximal insulin-stimulated (100 μU $ml^{-1}$) 2-deoxyglucose uptake in isolated and incubated soleus and/or EDL muscle from chow- (W) and HFD-fed (X) WT and mdKO male mice as well as chow- (Y) and HFD-fed (Z) WT and imdKO female mice.

(AA) AMP/ATP ratio in isolated and incubated soleus and EDL muscle from lean chow-fed female imdKO mice and WT littermates stimulated with metformin as described above.

(AB and AC) AMPK activity of the α2β2γ1, α1βγ1 and α2β2γ3 complexes in isolated and incubated soleus (AB) and EDL (AC) muscle from chow-fed male mdKO mice and WT littermates stimulated with metformin as described above.

(AD) Representative immunoblots of AMPKα Thr172, ACC Ser212, and TBC1D1 Ser231 phosphorylation as well as AMPKα2, ACC, TBC1D1, GLUT4 and HKII protein expression in isolated and incubated soleus and EDL muscle from chow-fed male mdKO mice and WT littermates stimulated with metformin and insulin as described above.

(AE) AMP/ATP ratio in isolated and incubated EDL muscle from HFD-fed female imdKO mice and WT littermates stimulated with metformin as described above.

(AF) AMPK activity of the α2β2γ1, α1βγ1 and α2β2γ3 complexes in isolated and incubated EDL muscle from HFD-fed male mdKO mice and WT littermates stimulated with metformin as described above.

(AG) Representative immunoblots of AMPKα Thr172, ACC Ser212, and TBC1D1 Ser231 phosphorylation as well as AMPKα2, ACC, TBC1D1, GLUT4 and HKII protein expression in isolated and incubated EDL muscle from DIO male mdKO mice and WT littermates stimulated with metformin and insulin as described above.

Data represent mean ± SEM. Data in panels A-D were analyzed using a homoscedastic two-tailed student’s t-test. Data in panels W and Y (soleus only) were analyzed by a mixed-effects model. Data in panels W (EDL only), X, Y (EDL only) and Z were analyzed using a three-way ANOVA with repeated measures. Data in the remaining panels were analyzed using a two-way ANOVA with (E, F, K, L, Q, and R) or without (G-J, M-P, S, AA-AC, AE, and AF) repeated measures. Sídák post hoc testing for multiple comparisons were used in all ANOVA analyses. * denotes $p<0.05$, ** denotes $p<0.01$, *** denotes $p<0.001$ and **** denotes $p<0.0001$. ME, main effect. INT, interaction.

**Figure S4**

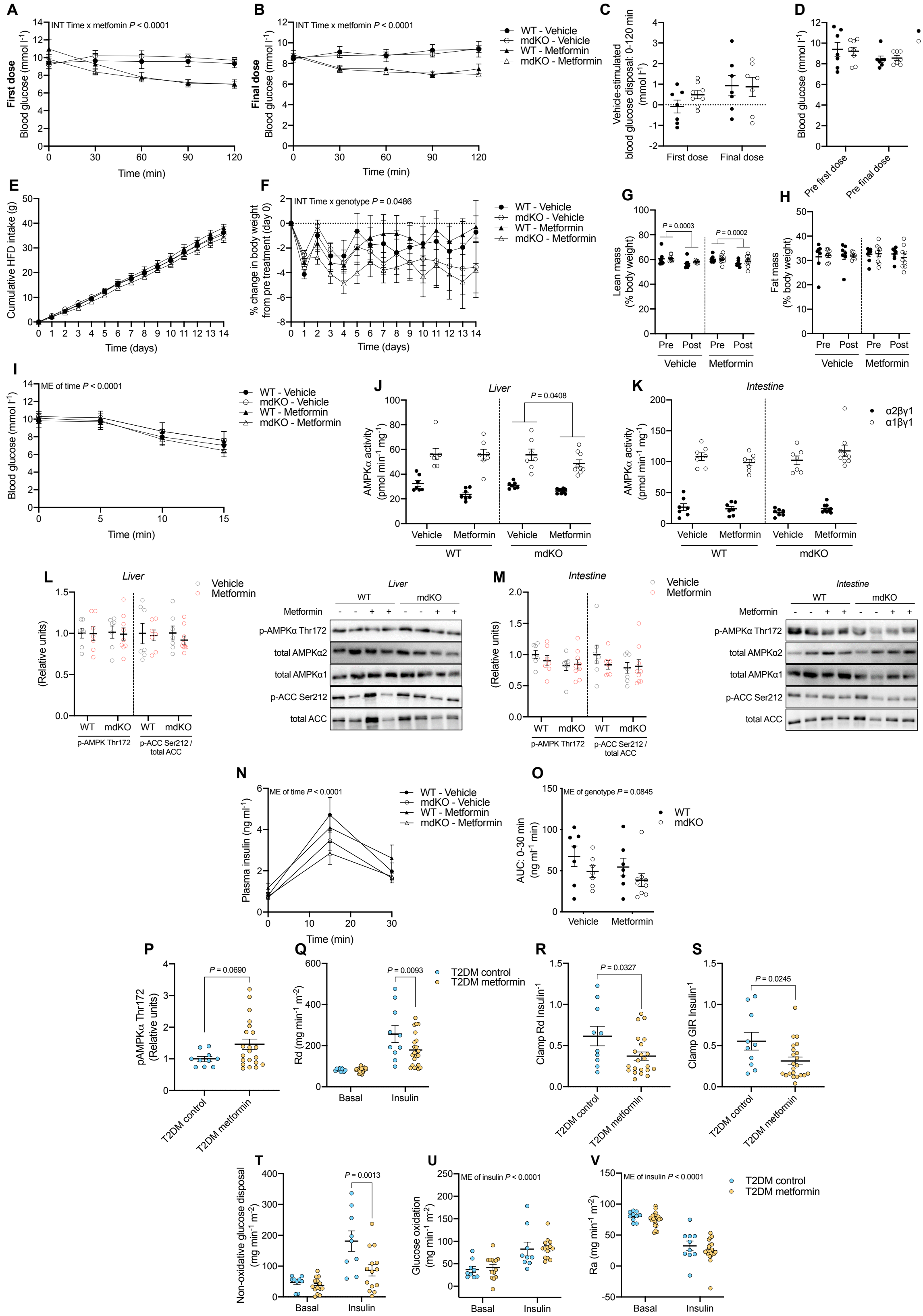


**Figure S4. Elevated AMPK signaling in human skeletal muscle following chronic metformin treatment is not associated with enhanced peripheral insulin sensitivity, Related to figure 4.**

(A and B) Effect of orally dosed metformin (250 mg $kg^{-1}$) vs. vehicle on blood glucose levels (mmol $l^{-1}$) in DIO AMPK mdKO mice and WT littermates. (A) initial dose (B) final dose of the chronic metformin treatment protocol (250 mg $kg^{-1}$ $day^{-1}$ for 14 days) (n = 7-9).

(C) Acute vehicle-stimulated changes in blood glucose levels (0-120 min) after the initial and final dose of the chronic metformin treatment period.

(D) Fed blood glucose levels prior to the first and final vehicle dose during the chronic metformin treatment period.

(E) Cumulative intake of HFD during the chronic metformin treatment period.

(F) Percent change in body weight from pretreatment during the chronic metformin treatment period.

(G and H) Body composition measured in DIO mdKO male mice and WT littermates before (Pre) and after (Post) the chronic metformin treatment period.

(I) Effect of retro-orbital administered insulin (1 Unit $kg^{-1}$) on blood glucose levels (mmol $l^{-1}$) in DIO AMPK mdKO mice and WT littermates after the chronic metformin treatment period.

(J-M) AMPK activity of the α2- and α1-associated complexes as well as phosphorylation of AMPKα Thr172 and ACC Ser212 in the liver (J and L) and intestine (K and M) of DIO AMPK mdKO mice and WT littermates after the chronic metformin treatment period. Right: representative immunoblots.

(N) Plasma insulin levels at time points 0, 15, and 30 min during a whole-body glucose tolerance test ~24 hours after the final dose of vehicle/metformin in DIO AMPK mdKO mice and WT littermates.

(O) Area under the curve (AUC) for plasma insulin during the glucose tolerance test (0-30 min).

(P) Phosphorylation of AMPKα Thr172 in muscle biopsy samples obtained from patients with type 2 diabetes mellitus treated with or without metformin and no other anti-diabetic medication. Metformin was withdrawn one week before the muscle biopsy sampling.

(Q-V) Measurements of Rd (Q), Clamp Rd $insulin^{-1}$ (R), Clamp GIR $insulin^{-1}$ (S), Non-oxidative glucose disposal (T), glucose oxidation (U), and Ra (V) during a hyperinsulinemic-euglycemic clamp in patients with type 2 diabetes mellitus treated with (yellow) or without (blue) metformin and no other anti-diabetic medication until one week before the clamp.

Data represent mean ± SEM. Data in panels P, R, and S were analyzed using a homoscedastic two-tailed student's t-test. Data in panels G and H were analyzed by a mixed-effects model. Data in panels A, B, E, F, I, and N were analyzed using a three-way ANOVA with repeated measures. Data in the remaining panels (C, D, J-M, O, Q, and T-V) were analyzed using a two-way ANOVA. Sídák post hoc testing for multiple comparisons were used in all ANOVA analyses. Rd, glucose rate of disappearance. GIR, glucose infusion rate. Ra, glucose rate of appearance. ME, main effect. INT, interaction.
